# Proteomic and Kinetic Analyses Reveal Discordant Apolipoprotein Turnover and Support a Revised Model of Human Lipoprotein(a) Metabolism

**DOI:** 10.64898/2026.09.04.749550

**Authors:** Tiffany Thomas, Anastasiya Matveyenko, Nelsa Matienzo, Michael Kissner, Santica Marcovina, Yihao Li, Haotian Wu, Stacia M Nicholson, Shiori Kuraoka, Masanori Aikawa, Sasha A. Singh, Gissette Reyes-Soffer

## Abstract

Lipoprotein(a) [Lp(a)] is a causal risk factor for atherosclerotic cardiovascular disease composed of apolipoprotein(a) [APO(a)] covalently linked to apolipoprotein B100 (APOB). Although plasma Lp(a) concentrations are largely genetically determined, the mechanisms governing Lp(a) metabolism after particle assembly remain poorly understood. We performed a cross sectional, observational study to define the Lp(a) proteome and determine the metabolic behavior of APO(a) and APOB within circulating Lp(a) particles using integrated proteomic, kinetic, and imaging approaches. Sixteen healthy adults underwent stable isotope tracer studies with ²H₃-L-leucine and ²H₅-glycerol. Lp(a) particles were isolated by APO(a)-specific immunoprecipitation for high-resolution liquid chromatography–mass spectrometry and kinetic analyses, and extracellular vesicles (EVs) were characterized by imaging flow cytometry and super-resolution microscopy. Proteomic analysis identified 92 proteins associated with immuno-isolated Lp(a), enriched in pathways related to immunity, coagulation, and atherogenesis. Lp(a)-APO(a) exhibited a mean fractional clearance rate of 0.04 pools/day, whereas Lp(a)-APOB cleared approximately sevenfold faster (0.25 pools/day), independent of plasma Lp(a) concentration or APO(a) isoform size. Both APO(a) and APOB were detected on circulating EVs, suggesting that EVs may contribute to post-secretory Lp(a) particle remodeling. These integrated human studies demonstrate marked discordance between APO(a) and APOB turnover within circulating Lp(a) particles, challenging the prevailing assumption that both apolipoproteins behave as a single metabolic unit after Lp(a) assembly and supporting a revised model of human Lp(a) metabolism.

## Introduction

Plasma protein concentrations are regulated by rates of production and clearance^1^ and in humans the processes can be quantified using stable isotopes and advances in mass spectrometry.^2^ Moreover, a protein’s residence time in circulation can and has been linked to disease mechanism.^3^ High plasma levels of Lipoprotein (a) [Lp(a)] is a causal risk factor for coronary artery disease, stroke, and aortic valve stenosis^4,5^, however critical questions remain in regards to assembly, secretion and clearance of Lp(a).^6,7^ Although the liver is the sole site of synthesis for the protein components of Lp(a), multiple receptors and intracellular trafficking routes appear to mediate Lp(a) synthesis, uptake and degradation.^7–9^ Rodent models lack Lp(a)^10^, and non-human primates^11^ have very low levels of Lp(a) levels and hepatic metabolism that differs from humans, limiting the translation of these animal models to a comprehensive understanding of Lp(a) regulation.

Lp(a) consists of two main protein components: an apolipoprotein B100 (APOB)–containing lipoprotein particle covalently and non-covalently bound to apolipoprotein(a) [APO(a)].^6,12^ APO(a) secretion can occur in the absence of normal APOB-containing lipoproteins, but assembly of mature circulating Lp(a) requires APOB-containing particles.^13^ APO(a) is a highly polymorphic glycoprotein encoded by the *LPA* gene^14^ which accounts for approximately 70-90% of the variability in plasma Lp(a) concentrations. Size heterogeneity of APO(a) arises from the variation in the number of Kringle IV type 2 (KIV-2) repeats, producing isoforms ranging from 1 to > 40 repeats.^14^ This variability underlies the well-described inverse relationship between Lp(a) plasma concentrations and isoform size.^4^

Human metabolic studies using stable isotopes are technically demanding, resources intensive, and require substantial volunteer commitment, limiting the number of studies performed. Interpretation and comparison of findings across studies are further complicated by differences in participant characteristics, feeding protocols, Lp(a) isolation methods, and participants metabolic state.^7,15^ These variations have resulted in inconsistent findings^7^, with reports suggesting either similar clearance rates for Lp(a)–APO(a) and Lp(a)–APOB compare to LDL-APOB^16^ or more rapid clearance of Lp(a)–APOB.^17^ Despite differences in methods, early work from DJ Rader et al ^18^ and others^7^ showed that small and large APO(a) isoforms had similar fractional catabolic rates (FCRs) and small APO(a) isoforms were produced more efficiently.

More recently, numerous studies have been performed describing the effects of novel APOB lowering therapies on Lp(a) metabolism ^19–22^. These studies were the first to establish that APO(a) production and APO(a) clearance are independently regulated and can be dissociated from LDL-APOB kinetics. In the current study, we performed stable isotope labeling studies in human participants with a wide range of Lp(a) concentrations, isolated intact plasma Lp(a) particles via immunoprecipitation (IP), followed by in-solution digestion, and high-resolution LC-MS analysis. Previous studies have used IP to isolate Lp(a), however these studies used gel electrophoresis to examine kinetics of the dominant APO(a) isoform. Our analysis of the intact Lp(a) particle allowed us to examine the proteomic profile (i.e., in solution digestion) and describe kinetic behavior of its two principal protein components, APO(a) and APOB. Finally, we revealed novel associations between circulating APO(a) and APOB particles with markers of extracellular vesicles (EVs), suggesting a novel mechanism underlying the prolonged residence time of Lp(a)-APO(a) relative to Lp(a)-APOB.

## Study Methods

### Study Population

Participants were recruited from the surrounding Tri-State area of Columbia University Irving Medical Center (CUIMC). All research related participant activities were performed at the Irving Institute for Clinical and Translational Research (IICTR), Clinical Research Resource (CRC) inpatient and outpatient facilities in New York, NY. The study enrolled and recruited subjects from 2019-2023.

Following consent, participants completed a screening visit that included a 12-hr fasting blood draw, medical history, physical exam (including electrocardiogram), a 24-hr dietary recall, and a taste test of the kinetic study liquid meal. Participants were enrolled within 4 weeks of screening and American Heart Association heart healthy dietary guidelines were provided.

Stable isotope studies were conducted in 16 individuals, and proteomic characterization was obtained in 14 of these individuals. For EV analysis, a randomly selected subgroup of participants with high (n = 4) and low (n = 3) plasma Lp(a) levels were selected, Detailed participant characteristics are provided in **Supplemental Table S1 and S2**.

### Sex as a biological variable

Sex was considered as a biological variable in study design through balanced enrollment of female and male participants (8 each). Because the study was not powered to detect sex-specific differences, all primary analyses were performed in the combined cohort. Sex was included as a covariate in multivariable association models to account for potential confounding, but sex-stratified analyses were not performed.

### Stable isotope administration

Participants were admitted to the inpatient research units of the CUIMC IICTR. On the morning of inpatient Day 0, blood samples were obtained in the outpatient research unit following a 12-hour overnight fast. Participants were allowed to leave the center and return at 5pm for admission. Before 8pm, dinner was subsequently provided by the staff of the IICTR Nutrition Research Unit after which subjects were made nothing by mouth (NPO). Beginning at 1am on Day 2, participants received liquid formula meals every 2 hours (16 feedings total) to achieve and maintain a steady nutritional state before, during, and post-administration of stable isotopes^63^. The composition of this liquid formula is 57% carbohydrate, 18% fat, and 25% protein.

At approximately 9am on day 2, baseline (0-hour) blood samples were collected for pre-stable-isotope measurement of various lipids and lipoproteins. Immediately thereafter, stable isotopes were administered for lipid and lipoprotein kinetics. Intravenous boluses of ^2^H3-L-leucine [10 μmol/kg body weight (BW)] and ^2^H5-glycerol (100 μmol/kg BW) were administered over a 10-minute period, followed by a constant intravenous infusion of ^2^H3-L-leucine (10 μmol/kg BW/hour) for 15 hours. Blood samples were collected at 0 (pre-bolus) and at various time-points post stable isotope administration. Participants were discharged on Day 3 after eating a heart healthy breakfast. Additional methodologic details are available in **Supplemental Method S1**.

### Plasma lipids and lipoproteins

Plasma lipid concentrations [total cholesterol (TC); triglycerides (TG), and HDL-C] were measured using the Integra 400plus (Roche). Plasma LDL-C(LDL-C) levels were estimated using NIH equation 2^64^. Plasma APOB concentrations were measured by a human enzyme-linked immunosorbent assay (ELISA; kit# 3715-1HP-2; Mabtech, Inc, Cincinnati, OH). ApoC3 was measured with a commercial ELISA (Abcam ab154131). Very low-density lipoprotein (VLDL) was isolated from timed plasma samples by sequential ultracentrifugation, as previously described^63^. We also measured oxidized phospholipids on APO(a) and APOB with a published method^65^.

### Lp(a) molar concentrations and apolipoprotein (a) size

Plasma Lp(a) concentrations were measured using an isoform-independent monoclonal antibody-based ELISA developed at the Northwest Lipid Metabolism and Diabetes Research Laboratory ^66,67^. APO(a) isoform size determination was performed by the same laboratory. Isoforms were isolated from 40 μl of plasma, and a standardized amount (100 ng) of APO(a) protein was loaded onto agarose gels, electrophoresed overnight at a final voltage of 123 V at 4°C, transferred to nitrocellulose membranes, immunoblotted, and imaged using a ChemiDoc MP Imaging System. APO(a) isoforms were resolved by size and identified by comparison with in-house standards consisting of individual plasma samples containing six apo(a) isoforms (38, 32, 24, 19, 15, and 12 KIV-2 repeats).

The relative expression of each isoform was quantified using Image Lab software, which calculated the proportional contribution of each isoform based on lane intensity profiles. Although estimation of isoform proportions depends on gel image quality and band intensity, the method exhibits inter-sample variability of less than 10%.

### Weighted allele size calculations

APO(a) isoform size is inversely correlated with plasma Lp(a) concentrations, with smaller isoforms generally predominating ^32^. To quantify the contribution of APO(a) isoforms to plasma Lp(a) levels, we calculated the weighted isoform size (*wIS*). The *wIS* was defined as the weighted average of the two APO(a) allele sizes, using the relative expression of each isoform determined from gel densitometry for each participant.

Example: If the two allele sizes are 20 and 30 KIV-2 repeats, with relative expression of 70% and 30%, respectively, the *wIS* is (0.7*20)+(0.3*30) = 23.

### Immunoprecipitation of Lp(a) from human plasma

Lp(a) was isolated from 500ul of frozen human plasma by immunoprecipitation (IP). Full methodological details are provided in the **Supplemental Method S2**.

### Characterization of Lp(a) proteome

Proteins identified with ≥3 unique peptides were included in the analysis. Relative protein abundances were visualized using normalized protein abundance values. Average protein abundances across time points were used to generate heat map. Additional methodological details are provided in the **Supplemental Method S3**. The mass spectrometry proteomics data have been deposited to the ProteomeXchange Consortium via the PRIDE partner repository with the dataset identifier PXD081142 and 10.6019/PXD081142

### Stable isotope enrichment

Gas chromatography mass spectrometry (GC-MS) was used to measure enrichment of ^2^H3-L-leucine and Ring-^13^C6-L-phenylalanine in VLDL-APOB100 using an Agilent 6890 GC and a 5973 MS operating in negative chemical ionization mode, following previously published protocols, expanded methods can be found in **Supplemental Method S3 and S4 and Supplemental Table S3, S4**.

### Stable isotope enrichment data modeling

APOB turnover (i.e. kinetics) in VLDL was determined using ^2^H3-L-leucine enrichment data obtained from the bolus injection and constant infusion phases. VLDL-APOB enrichment data were fitted using a multicompartmental model implemented in Poolfit ^68^ which solves the differential equations in closed form and computes parameter estimates and sensitivities as sums of exponentials.

Lp(a)-APO(a) and Lp(a)-APOB FCR were calculated by fitting leucine enrichment data from peptide specific enrichment measurements using a single-pool model, with the precursor enrichment defined by the VLDL APOB enrichment plateau (see above). The plateau is typically reached during the 15-hr sampling period and was estimated using our established model for APOB metabolism. The Lp(a)-APO(a) production rate (PR; nmol/kg/day) was calculated as the product of APO(a) FCR (pools/day) and the Lp(a) concentration (nmol/L), multiplied by the estimated plasma volume (0.045 L/kg). The Lp(a)-APOB PR was calculated using the APOB concentration, expressed as percentage of total APOB associated with Lp(a) and the molecular mass of APOB (550kDa).

### Plasma isolation of extracellular vesicles and characterization using imaging flow cytometry

Total plasma EVs were isolated via size exclusion chromatography (SEC) [SmartSEC™ Mini EV Isolation System (System Biosciences, SSEC100A-1)] . Importantly, we isolated an EV enriched fraction (Fraction 1 of 3 from the kits). The data presented are all from this fraction. Canonical EV markers (CD9, CD63, CD81) and their association to APO(a) and APOB were assessed using a previously published flow cytometry method.^69^ Representative imaging flow cytometry images acquired on an ImageStreamX Mk II and analyzed using IDEAS v6.2 software, **Supplemental Figure S5.** Submicron particles were selected by gating on objects with side scatter measurements lower than those of 1 μm Amnis SpeedBead particles. Channels shown include LPA (AF647), ApoB (PE), CD9 (PE-Cy7), CD63 (BV421), CD81 (PE-Dazzle), and the composite image. Scale bar: 7 μm.

Super resolution nanoimaging of the EVs was also performed^70^ in a subset (N=3) of our subjects, confirming the associations found via flow cytometry imaging**, Supplemental Figure S6c and S7.** In addition, we performed negative-stain transmission electron microscopy (TEM) showing the morphology and size range of isolated extracellular vesicles (EVs) and immunogold TEM demonstrating EV, APO(a) marker labeling on EVs, with LPA4, **Supplemental Figure S6 (a) and (b)**. Additional methodological details are provided in **Supplemental Method S5. Supplemental Figure S5 and S6** highlight the EVs characterization summary.

## Statistical analysis

Associations between protein abundances, plasma measurements [Lp(a) and wIS], kinetic parameters (PR and FCR), and EV particle concentrations were assessed using ordinary linear regression models adjusted by age, sex, and race. Protein abundances and EV particle concentration were median scaled prior to analysis. Lp(a) concentrations were natural-log transformed to account for right-skewed distributions (results based on raw value are also included in supplemental data for reference). P values for coefficients of interest were adjusted for multiple testing using the Benjamini-Hochberg false discovery rate (FDR) procedure. Statistical significance was defined as nominal P < 0.05 and trends were defined as P < 0.10 for the small sample size. While the FDR adjusted P < 0.05 are also reported for reference. All analyses were performed using the Statsmodels (v0.13.1) with solver SciPy (v1.7.2) in Python (v3.8.12). Some preprocessing pipelines are implemented in R (v4.4.1).

The data supporting the findings of this study are available within the article and its supplemental materials. Source data for all graphs and summary statistics are provided in the supporting data. Additional data is available from the corresponding author upon reasonable request.

### Study Approval

All study procedures were approved by the Columbia University Medical Center Institutional Review Board (IRB AAAR9251) and the IICTR, and all participants provided written informed consent prior to enrollment.

### Data availability

Additional data and analytical codes not included in study methods and supplemental data will be provided upon reasonable request.

## Results

### Study population

We enrolled sixteen healthy volunteers (8 females and 8 males) into a cross-sectional observational mechanistic study. Baseline characteristics are summarized in **Table 1**, and individual-level data are provided in **Supplemental Tables S1 and S2.** The mean age of the cohort was 46 ± 14 years. Female and male participants were equally represented to account for sex as a biological variable. Because the study was not powered to detect sex-specific differences, all primary analyses were performed in the combined cohort, with sex included as a covariate in multivariable association analyses. The cohort self-reported race was diverse and included non-Hispanic White, Black, and Caribbean Hispanic participants. The mean body mass index (BMI) of the cohort was 28.59 kg/m² with no history of diabetes or hypertension and normal blood chemistry profiles.

**Table 1.**
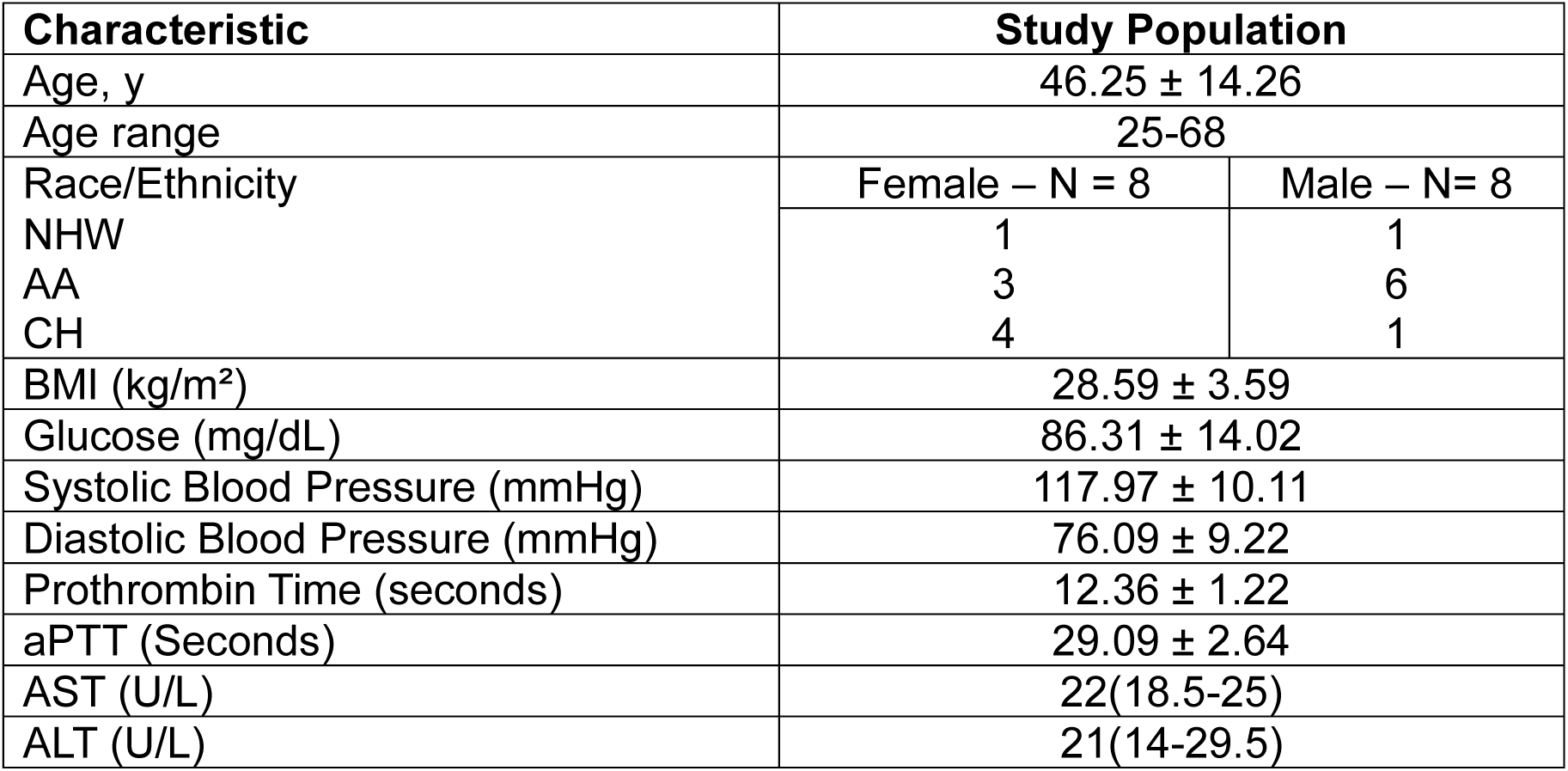
Baseline Characteristics. **Legend:** Baseline characteristics for study cohort. Race/Ethnicity are self-reported. NHW: non-Hispanic White, B: Blacks, CH: Caribbean Hispanic; aPTT: Activated partial thromboplastin time, AST: Aspartate aminotransferase, ALT: alanine aminotransferase; ± represents mean and standard deviation for normally distributed variables; ( ) represents median and interquartile range.

### Lipid and apolipoprotein profiles

Baseline plasma lipids and apolipoproteins are summarized in **Table 2**. Most lipid parameters were within normal reference ranges, except for total cholesterol (207.31 ± 44.93 mg/dL) and LDL-C (Mean 129 ± 43.29 mg/dL) which were modestly elevated in 8 out of the 16 subjects. HDL-C (59.38 ± 16.72 mg/dL) and TG [ 87.5 mg/dL, interquartile range (IQR) 67-145.5 mg/dL] were within normal limits for the majority of subjects (15/16 and 12/16 for HDL-H and TG respectively). The median Lp(a) concentration was 58.96 nmol/L (IQR 25.51-133.35). APOC-III concentration was also within normal reference range with one subject in the borderline range (116.1mg/L). As oxidized phospholipids have been shown to drive Lp(a) atherogenicity we wanted to measure the levels in this population. Median OxPL-APO(a) and OxPL-APOB concentrations were modestly higher than those reported in healthy reference populations, consistent with the moderately high Lp(a) concentrations in our cohort.

**Table 2.**
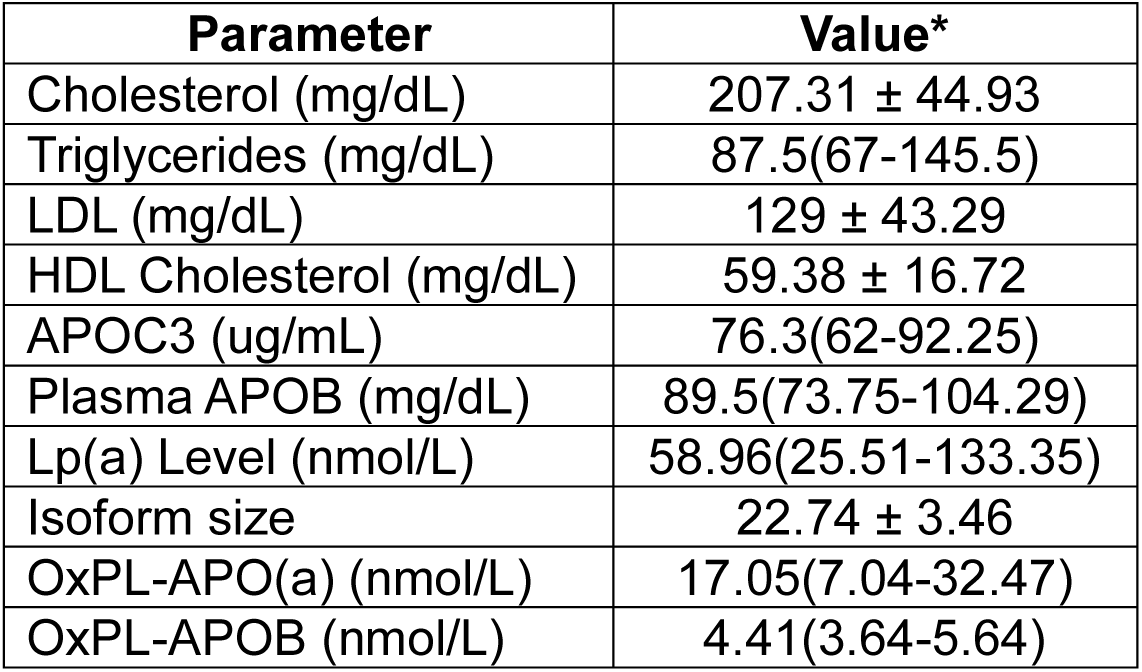
Baseline lipid, apolipoprotein, and oxidized phospholipid measurements. **Legend:** LDL: low density lipoprotein, HDL: high density lipoprotein, APO: apolipoprotein, APOB: apolipoprotein B100, Lp(a): lipoprotein(a), OxPL: oxidized phospholipid. Isoform sizes are presented as calculated weighted isoform size levels. * (±) represents mean and standard deviation for normally distributed variables; ( ) represents median and interquartile range.

### Lp(a) proteome

Proteome analysis identified a total of 92 distinct proteins in immuno-isolated Lp(a) particles including APO(a) and APOB (**Supplemental Figure S1, S2 and Supplemental Table S3**). In line with our recent publication^23^, key proteins that highlight the role of Lp(a) in lipid metabolism pathways but also as a carrier of coagulation, complement and inflammatory signals. Exploratory proteomic analyses identified several proteins nominally associated with plasma Lp(a) concentration, with the majority clustering within biologically related pathways of coagulation, complement activation, and hepatic protein transport, **Table 3**. Higher plasma Lp(a) concentrations were positively associated with proteins involved in coagulation and thrombosis, including kininogen-1 (β = 1.197, *P* = 0.0069) and antithrombin III (β = 0.435, *P* = 0.0157), whereas thrombospondin-1 demonstrated an inverse association (β = –0.191, *P* = 0.0276). Within innate immune pathways, complement C1s was positively associated with plasma Lp(a) concentration (β = 0.720, *P* = 0.0274), while complement C4-B was inversely associated with weighted APO(a) isoform size (β = –2.292, *P* = 0.0100). Additional positive associations with plasma Lp(a) concentration were observed for proteins involved in hepatic transport and acute-phase responses, including serotransferrin, albumin, α2-macroglobulin, transthyretin, and α1-acid glycoprotein-1 (all nominal *P* < 0.05). None of the associations remained statistically significant following false discovery rate correction, **Supplemental Table S5.**

**Table 3.** Proteins nominally associated with plasma Lp(a) concentration and weighted apo(a) isoform size grouped by biological pathway. **Legend: Exploratory associations between the Lp(a) proteome, plasma Lp(a) concentration, and apo(a) isoform size.** Linear regression analyses were performed to identify proteins associated with plasma Lp(a) concentration (log-transformed) and weighted APO(a) isoform size. Proteins are grouped according to their predominant biological function to facilitate biological interpretation. Positive β coefficients indicate a direct association between protein abundance and the outcome, whereas negative β coefficients indicate an inverse association. Nominal (*P* < 0.05) and false discovery rate (FDR)-adjusted P values are shown. Although none of the associations remained statistically significant after FDR correction, the proteins clustered within biologically coherent pathways related to coagulation, complement activation, and hepatic protein transport, supporting their use as hypothesis-generating candidates for future mechanistic investigation. β, regression coefficient; FDR, false discovery rate; Lp(a), lipoprotein(a).

| Biological Pathway | Protein | Outcome | $\beta$<br>Coefficient | P<br>Value | FDR-adjusted P<br>Value |
| --- | --- | --- | --- | --- | --- |
| **Coagulation and thrombosis** | Kininogen-1 | Plasma Lp(a) concentration | 1.197 | 0.0069 | 0.220 |
|  | Antithrombin III | Plasma Lp(a) concentration | 0.435 | 0.0157 | 0.241 |
|  | Thrombospondin-1 | Plasma Lp(a) concentration | -0.191 | 0.0276 | 0.283 |
| **Complement and innate immunity** | Complement C1s | Plasma Lp(a) concentration | 0.720 | 0.0274 | 0.283 |
|  | Complement C4-B | Weighted apo(a) isoform size | -2.292 | 0.0100 | 0.917 |
| **Hepatic transport and acute-phase response** | Serotransferrin | Plasma Lp(a) concentration | 0.383 | 0.0068 | 0.220 |
|  | Albumin | Plasma Lp(a) concentration | 0.441 | 0.0078 | 0.220 |
| | α2-Macroglobulin | Plasma Lp(a) concentration | 0.256 | 0.0096 | 0.220 |
|  | Transthyretin | Plasma Lp(a) concentration | 0.978 | 0.0138 | 0.241 |
| | α1-Acid Glycoprotein 1 | Plasma Lp(a) concentration | 0.329 | 0.0240 | 0.283 |

### Fractional clearance and production of Lp(a)-APO(a) and Lp(a)-APOB

We used a linear model to fit leucine enrichment data from peptide-specific measurements, **Supplemental Figure S3 APO(a) and S4 (APOB).** The mean Lp(a)-APO(a) FCR was 0.04 pools/d (range: 0.01 to 0.17), whereas Lp(a)-APOB FCR was approximately seven-fold higher with a mean of 0.25 pools/d (range: 0.08 to 0.44). Production rates (PRs) were calculated using the plasma pool size of Lp(a) and APOB, yielding a mean Lp(a)-APO(a) PR of 0.12 nmol/kg/d (range: 0.01 to 0.39) and Lp(a)-APOB PR of 0.81 nmol/kg/d (range: 0.12 to 2.51), **Table 4**, **Figure 1**. In this cohort, both APO(a) and APOB PRs were positively associated with log transformed Lp(a) concentrations (FDR-adjusted P=0.007). The APO(a) isoform size (wIS) showed an inverse trend associated with Lp(a)-APOB FCR and PR (P= 0.052, 0.054, respectively), **Supplemental Table S6**.

**Figure 1.**
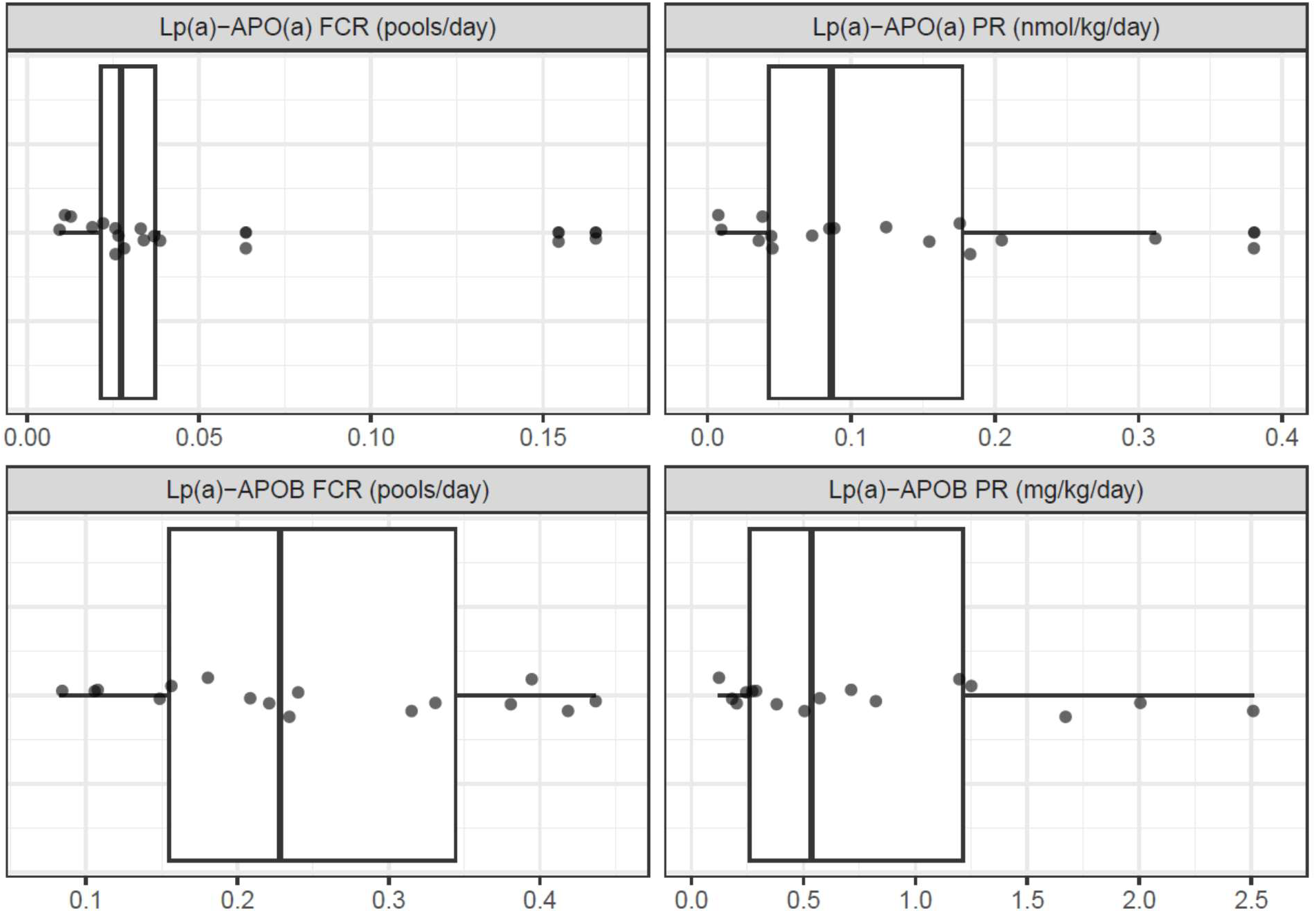
**Distribution of Lp(a) apolipoprotein kinetic parameters in healthy adults**. Boxplots summarize the distribution of each kinetic parameter. The box represents the interquartile range (IQR; 25th–75th percentiles), and the horizontal line within the box indicates the median. Whiskers extend to the most extreme observations within 1.5 × IQR of the lower (Q1) and upper (Q3) quartiles. Individual points beyond the whiskers are displayed as outliers, while all observations are shown as overlaid data points. The distributions are generally right-skewed. Note that the x-axis scales differ across panels.

**Table 4.** Fractional catabolic rates and production rates for Lp(a)–APO(a) and Lp(a)–APOB. **Legend:** Fractional catabolic rates (FCR) and production rates (PR). APO: apolipoproteins. The FCR’s and PR’s were derived from stable isotope enrichment curves in 16 healthy participants. Data are presented as minimum, mean, median, interquartile range (IQR), and maximum values. FCR is expressed in pools per day, and PR is expressed as mg/kg/day.

|  | **Min** | **Mean** | **Median** | **IQR** | **Max** |
| --- | --- | --- | --- | --- | --- |
| APO(a) FCR<br>pools/day | 0.009 | 0.044 | 0.027 | 0.021-0.037 | 0.165 |
| APOB FCR<br>Pools/day | 0.082 | 0.248 | 0.228 | 0.155-0.344 | 0.437 |
| APO(a) PR<br>nmol/kg/day | 0.007 | 0.123 | 0.086 | 0.043-0.178 | 0.381 |
| APOB PR<br>mg/kg/day | 0.118 | 0.808 | 0.536 | 0.26-1.213 | 2.512 |

Among the 92 proteins identified, exploratory analyses identified several proteins nominally associated with Lp(a) kinetic parameters, **Table 5**. Proteins positively associated with APO(a) fractional clearance rate (FCR) and/or production rate (PR) included prenylcysteine oxidase 1 (PCYOX1), apolipoprotein D, complement C3 (C3), and clusterin (ApoJ). In contrast, cholesteryl ester transfer protein (CETP), phospholipid transfer protein (PLTP), complement C1q subunits B and C, and heparin cofactor II were inversely associated with APO(a) kinetics. ApoB kinetics demonstrated fewer associations, with APOC-II positively associated with APOB FCR. Several proteins, including PCYOX1, apolipoprotein D, and C3, were associated with both APO(a) and APOB kinetic parameters, **Supplemental Table S7**.

**Table 5.** Proteins nominally associated with Lp(a) kinetic parameters and biological pathways. Legend: Regression analyses performed to identify proteins associated with Lp(a)-APO(a) and Lp(a)- APOB fractional clearance rates (FCRs) and production rates (PRs). Numerous proteins demonstrated nominal associations (*P* < 0.05) with one or more kinetic parameters, including the direction of association, principal kinetic endpoint(s) and biological function. Included: regression coefficients (β), nominal *P* values, and false discovery rate (FDR)-adjusted *P* values. APO(a), apolipoprotein(a); APOB, apolipoprotein B100; CETP, cholesteryl ester transfer protein; PLTP, phospholipid transfer protein; FCR, fractional clearance rate; PR, production rate; HDL, high-density lipoprotein; Lp(a), lipoprotein(a).

| **Protein** | **Primary association** | **$\beta$** | **Nominal P** | **Biological relevance** |
| --- | --- | --- | --- | --- |
| **Prenylcysteine oxidase 1 (PCYOX1)** | APO(a) FCR<br>APO(a) PR<br>APOB PR | ↑ | 0.004<br>0.025<br>0.021 | Oxidative lipid metabolism; pro-atherogenic |
| **Apolipoprotein D** | APO(a) FCR<br>APO(a) PR<br>APOB PR | ↑ | 0.021<br>0.024<br>0.018 | Lipid transport and oxidative stress |
| **Complement C3** | APO(a) FCR<br>APO(a) PR<br>APOB PR | ↑ | 0.047<br>0.004<br>0.009 | Complement activation; vascular inflammation |
| **CETP** | APO(a) FCR<br>APO(a) PR | ↓ | 0.007<br>0.030 | Lipid exchange between lipoproteins |
| **PLTP** | APO(a) FCR<br>APO(a) PR | ↓ | 0.006<br>0.002 | HDL remodeling; phospholipid transfer |
| **Clusterin (ApoJ)** | APO(a) FCR | ↑ | 0.003 | Chaperone; HDL/Lp(a) associated |
| **Complement C1q B/C** | APO(a) FCR | ↓ | 0.001/0.017 | Complement activation |
| **APOC-II** | APOB FCR | ↑ | 0.014 | Triglyceride metabolism |

### Associations of APO(a) and APOB with EVs

Based on findings from our stable isotope studies, and the long half-life of APO(a) particles, we hypothesized that apoproteins particles may undergo recycling^7,24^ and recent studies have reported associations between LDL particles and EVs.^25, 26–28^ To explore this possibility, we conducted an exploratory analysis of EVs with APO(a) and APOB in seven out of the 16 participants who completed stable isotope studies, including four with high Lp(a) concentrations and three with low Lp(a) concentrations. Participant characteristics for this small study are highlighted in **Supplemental Table S1**. The ImageStream imaging flow cytometry workflow used to distinguish free from EV-associated APO(a) and APOB particles, along with representative images, is provided in **Supplemental Figure S5**. Representative flow cytometry gating strategy can be found on **left panel** (a and b) and **right panel** (c) images demonstrate presence of APO(a) and APOB overlying EV structures. The EV structures are detected by presence of tetraspanins (CD9, CD63, CD81). These overlapping fluorescence signals within the same diffraction-limited particle, visualized in the composite images, support physical association of APO(a) and APOB on the same EV rather than coincidence detection of separate particles. Importantly, our EV enriched fractions did contain free (not associated to tetraspanins) apoproteins. These data are consistent with a model in which APOB/APO(a) are associated with tetraspanin-positive EVs, thereby serving as an optional platform for their transport. After FDR-adjustment, no associations were observed between EV associated apoproteins and plasma Lp(a) concentrations or isoform size, **Supplemental Table S8.** However, we observed a nominal positive trend between Lp(a)-APO(a) FCR and EV-associated APO(a), as well as a significant association between free APOB and Lp(a)-APOB FCR, **Table 6, Supplemental Table S9.**

**Table 6.** Associations between extracellular vesicle–associated apolipoproteins and Lp(a) kinetic parameters. **Legend:** Regression analyses to examine relationships between circulating extracellular vesicle (EV)- associated apolipoproteins and Lp(a) kinetic parameters. β coefficients represent the direction and magnitude of association between each EV marker and the corresponding kinetic trait. Free APOB demonstrated a significant positive association with Lp(a)-APOB fractional clearance rate (FCR) after false discovery rate (FDR) correction. EV-bound APO(a), EV-bound APOB, and EVs containing both APO(a) and APOB demonstrated trend-level associations with Lp(a)-APO(a) kinetic parameters but did not reach statistical significance following FDR correction. These findings support a potential role for extracellular vesicles in Lp(a) metabolism and identify candidate pathways linking EV-associated apolipoproteins with apo(a) production and clearance. apo(a), apolipoprotein(a); APOB, apolipoprotein B100; EV, extracellular vesicle; FCR, fractional clearance rate; FDR, false discovery rate; PR, production rate.

| Lp(a) Kinetic Trait | EV Marker | $\beta$ | FDR-adjusted P Value | Interpretation |
| --- | --- | --- | --- | --- |
| Lp(a)-APOB FCR | Free APOB | 0.24 | 0.004 | Faster APOB turnover |
| Lp(a)-APO(a) FCR | EV-bound APO(a) | 0.02 | 0.098 | Trend toward increased APO(a) clearance |
| Lp(a)-APO(a) PR | EV-bound APOB | −0.07 | 0.103 | Trend toward reduced APO(a) production |
| Lp(a)-APO(a) PR | EV-bound APO(a)+APOB | −0.05 | 0.103 | Suggests EV-mediated regulation of APO(a) production |

## Discussion

Our findings support a revised model in which APO(a) and APOB do not remain metabolically linked after secretion. Instead, APOB enters the canonical LDL metabolic pathway, whereas APO(a) persists within a slowly turning-over compartment that may undergo recycling, delayed reassociation with newly secreted APOB-containing particles, or trafficking through extracellular vesicles. This model reconciles our kinetic observations with previous human intervention studies and provides a mechanistic framework for understanding the effects of emerging Lp(a)-lowering therapies.

*Lp(a) proteome and mechanistic insights.* Our proteomic analysis of Lp(a) identified enrichment of pathways previously implicated in cardiovascular disease pathogenesis.^29,30^ As expected classical apoproteins were identified such as APOB, APOCII, APOCIII. In addition, proteins involved in innate immunity and complement activation (complement C1s), coagulation (antithrombin III), and inflammatory responses (α1-acid glycoprotein-1) increased in those with higher plasma Lp(a), consistent with the well-established links between Lp(a), thrombosis, and vascular inflammation.

Likewise, associations with serotransferrin, transthyretin, and albumin may reflect broader alterations in plasma protein transport or hepatic protein secretion accompanying elevated Lp(a) concentrations. The inverse association between thrombospondin-1 and Lp(a) is intriguing given its role in extracellular matrix remodeling and platelet activation but requires independent validation. Similarly, the negative association between complement C4-B and APO(a) isoform size raises the possibility that immune-related pathways differ according to APO(a) isoform, although this observation should be interpreted cautiously.

Although these exploratory associations require validation, they collectively implicate biologically relevant pathways involved in lipoprotein remodeling, oxidative stress, innate immunity, and coagulation. Importantly, these associations were observed predominantly with Lp(a)-APO(a), rather than Lp(a)-APOB, kinetic parameters, despite both proteins residing on the same Lp(a) particle.

Together with the marked discordance in APO(a) and APOB turnover, these findings support the concept that APO(a) metabolism is governed by biological pathways distinct from those regulating APOB-containing lipoproteins. The convergence of proteins with established roles in lipid remodeling (CETP, PLTP, APOD), oxidative biology (PCYOX1), complement activation (C3, C1q), and thrombosis (α2-antiplasmin, heparin cofactor II) provides a mechanistic framework for future studies examining the regulation of Lp(a) assembly, remodeling, and clearance.

These findings raise the possibility that Lp(a) is accompanied not only by a greater number of circulating particles but also by qualitative changes in particle composition that enhance its biological activity. Such compositional remodeling could augment the inflammatory, immune, and prothrombotic properties of Lp(a), providing a mechanism by which higher Lp(a) concentrations disproportionately amplify cardiovascular risk. Collectively, our data supports the emerging concept that Lp(a) is not merely a cholesterol-carrying lipoprotein, but a dynamic carrier of bioactive proteins involved in vascular inflammation, coagulation, and immune regulation.

*Lp(a) apo(a) and apoB100 metabolism.* The metabolic fate of secreted Lp(a) particles has been investigated using diverse methodological and experimental approaches and has been extensively reviewed. ^6,7^ Several, although not all, stable isotope studies in humans observed similar FCR values for Lp(a)-APO(a) and Lp(a)-APOB supporting the concept that Lp(a) particles are assembled intracellularly and secreted with both APO(a) and APOB. However, prior kinetic studies relied on protein isolation via gel electrophoresis entailing proteolysis of select molecular weight bands, whereas our study used in solution digestion of entire immunopurified Lp(a). This distinction is important because a single isoform can have different kinetics from the pooled APO(a) signal. In-gel proteolysis may isolate a specific isoform, whereas in-solution digestion reflects the combined APO(a) population.

Early work by Rader and colleagues demonstrated that APO(a) isoform size regulates Lp(a) production.^18^ Subsequent studies, including those from our group, confirmed these early observations and further demonstrated that plasma Lp(a) concentrations are determined predominantly by APO(a) production rather than fractional catabolic rate in individuals expressing small APO(a) isoforms.^31,32^ In contrast, in individuals with larger APO(a) isoforms, both APO(a) production and clearance were associated with plasma Lp(a) concentrations. Chan et al. further reported that both APO(a) PR and APO(a) isoform size independently predict Lp(a) levels in multivariable models, suggesting that factors beyond APO(a) isoform size contribute to APO(a) production.^22^ Despite important methodological differences in Lp(a) isolation and kinetic modeling across studies, our finding that APO(a) exhibits substantially slower clearance than APOB is consistent with the independent observations of Jenner et al., Diffenderfer et al and others.^16,33,34^ Furthermore, the Lp(a)-APOB fractional catabolic rate observed in our study closely approximates previously reported values for LDL-APOB turnover^35^,supporting the concept that APOB metabolism remains largely conserved after incorporation into the Lp(a) particle.^32,33^ In contrast, Lp(a)-APO(a) FCR resembled that of long-lived plasma proteins such as albumin, which are maintained at steady state concentrations and undergo recycling.^36,37^

The magnitude of this kinetic divergence was unexpected. We observed markedly different turnover rates for Lp(a)-APO(a) (estimated half-life ≈17 days) vs. and Lp(a)-APOB (estimated half-life ≈2.8 days). Because we immunoprecipitated intact circulating Lp(a), in which each APO(a) molecule is covalently linked to a single APOB molecule, both tracers should have exhibited similar kinetics if they remained associated throughout their metabolic lifetime. Instead, their strikingly different turnover rates indicate that APO(a) and APOB become metabolically uncoupled after secretion.

This observation represents the major mechanistic finding of our study. It supports a working model in which APO(a) resides within a metabolically distinct, slowly equilibrating precursor pool that is not kinetically coupled to the APOB pool measured within intact Lp(a). In this model, APO(a) may remain in circulation for prolonged periods before associating with newly synthesized APOB-containing particles or with APOB particles already present in plasma. Conversely, APOB likely re-enters the conventional LDL metabolic pathway, including LDL receptor-mediated clearance, whereas APO(a) follows a slower pathway^38^ involving recycling onto newly available APOB particles and/or association with extracellular vesicles, thereby explaining its prolonged plasma residence time.

Although *in vivo* human studies cannot directly resolve the intracellular mechanisms governing Lp(a) assembly, experimental studies in hepatocytes and animal models provide important mechanistic support. Non-covalent APO(a)-APOB interactions are proposed to precede extracellular disulfide bond formation, leading to the concept that APOB functions as a “chaperone” for APO(a) secretion^12^. Consistent with this model, patients with abetalipoproteinemia, who lack circulating APOB-containing lipoproteins, secrete APO(a) but fail to form circulating Lp(a) particles, demonstrating that APOB is required for Lp(a) assembly^13^.Similarly, patients with microsomal triglyceride transfer protein (MTP) deficiency^39^, in whom APOB-containing lipoprotein secretion is severely impaired, have extremely low circulating Lp(a) concentrations, further supporting the requirement for APOB secretion in Lp(a) assembly.

*Additional support comes from human intervention studies.* Therapeutic reduction of APO(a) synthesis using antisense oligonucleotides lowers circulating APO(a) by up to 90%^40^ without comparable reductions in LDL-APOB, demonstrating that APO(a) synthesis can be selectively manipulated independently of bulk APOB metabolism. Likewise, APO(a) disruptor therapies inhibit the interaction between APO(a) and APOB during particle assembly^41,42^, indicating that APO(a) synthesis and APO(a)-APOB assembly are mechanistically separable processes.

Further support comes from the MAESTRO-NASH program^43^, in which treatment with the thyroid hormone receptor-β agonist resmetirom produced a greater reduction in Lp(a) than in APOB despite marked improvements in hepatic steatosis and lipid metabolism. These findings are difficult to reconcile with a single homogeneous APOB secretory pool. Instead, they suggest that hepatic lipid remodeling preferentially alters the subset of newly synthesized APOB particles available for Lp(a) assembly rather than uniformly reducing total APOB production.

This interpretation is consistent with evidence that although most circulating LDL-APOB arises through VLDL remodeling, approximately 10% is secreted directly by the liver^44^, potentially providing a triglyceride-poor APOB substrate for APO(a) association. Moreover, the relatively small number of circulating Lp(a) particles compared with LDL-APOB supports the existence of a specialized APOB subpopulation dedicated to Lp(a) formation.

In contrast to the compositional changes associated with circulating Lp(a) concentrations, none of the protein associations with FCR or PR remained significant after false discover rate (FDR) correction. This suggesting that Lp(a) particle turnover may be influenced by a limited subset of biologically relevant interactions rather than broad proteomic remodeling; nevertheless, the nominal associations provide several hypothesis-generating leads. These associations clustered predominantly within three biological domains. First, associations with proteins involved in lipoprotein remodeling, including CETP, PLTP^45^, APOD^46^, and APOC-II, suggest that differences in Lp(a) kinetics may relate to exchange of surface lipids and apolipoproteins rather than simply particle concentration. Second, associations with complement proteins C3, C1qB, C1qC implicate innate immune pathways. Third, the association with prenylcysteine oxidase (PCYOX1) highlights a potential role for oxidative biology. PCYPX1, which has been identified in the proteome in a few previous studies. It generates hydrogen peroxide, promotes LDL oxidation, and contributes to vascular inflammation. Its nominal association with both APO(a) and APOB production makes it one of the strongest mechanistic candidates for future studies. These findings are consistent with emerging evidence demonstrating crosstalk between coagulation, extracellular matrix remodeling, and lipoprotein metabolism^47^ and further support a role for Lp(a) in plaque development and progression^48^. Notably, most nominal associations involved APO(a) FCR or PR, whereas APOB kinetics show only a small number of proteins including PCYOX1, APOD, C3, APOC-II, and Kininogen-1.

*Association with extracellular vesicles.* To further investigate the markedly different residence times of APO(a) and APOB within circulating Lp(a), we examined their association with plasma extracellular vesicles (EVs). Although EV preparations were historically considered to be contaminated by lipoproteins because of their overlapping size and density, recent studies have demonstrated that LDL forms a stable biomolecular corona on EV surfaces^49^ and liver derived EVs contain APOB. ^28^ Emerging evidence further indicates that lipoprotein-EV interactions influence extracellular and intracellular transport, alter uptake by recipient cells, and may facilitate endothelial transcytosis through lipoprotein trafficking pathways. ^26,50–52^ Proteomic studies have likewise identified multiple apolipoproteins within EV preparations.^53^

Our imaging flow cytometry and super-resolution nanoscopy provide direct evidence that both APO(a) and APOB are closely associated with circulating EVs. While these observations do not establish the functional consequences of this interaction, they identify EVs as a previously underappreciated compartment for Lp(a)-associated apolipoproteins that may contribute to their transport, compartmentalization, or prolonged plasma residence. This finding is particularly noteworthy considering our kinetic data demonstrating markedly slower turnover of APO(a) than APOB, raising the possibility that EV association represents one mechanism underlying their divergent metabolic behavior.

The concept of EV-mediated apolipoprotein trafficking is biologically plausible. Classical studies established that exchangeable apolipoproteins, particularly APOE and the APOCs, readily dissociate from lipoprotein surfaces and redistribute among lipoprotein particles through aqueous diffusion. ^54 55,56^ Although APO(a) is covalently linked to APOB within mature Lp(a), our findings raise the possibility that APO(a)-or APOB-containing particles, or their associated protein complexes, may similarly interact with EV surfaces, facilitating redistribution, recycling, or transport to specific tissues. This concept is further supported by growing evidence that apolipoproteins participate in EV assembly, trafficking, and biological function.^49^

Our observations also suggest a potential mechanism for extracellular Lp(a) assembly. EVs have been shown to carry protein disulfide isomerase (PDI) family members and other thiol oxidoreductases capable of catalyzing disulfide bond formation, reduction, and isomerization in the extracellular environment^57^. Extracellular PDI likewise mediates thiol-disulfide exchange reactions on cell surfaces and within the vasculature, demonstrating that extracellular redox-dependent protein remodeling is biologically feasible. Given that the oxidase-like activity originally described by Koschinsky and colleagues^58^ remains molecularly unidentified, hepatocyte-derived EVs could provide a localized catalytic surface that concentrates APO(a) and APOB while simultaneously promoting disulfide bond formation during extracellular Lp(a) assembly. Although direct evidence for EV-mediated Lp(a) assembly is not yet available, this hypothesis provides a plausible mechanistic explanation for the extracellular, proteinaceous catalytic activity previously observed in conditioned media and offers a testable framework for future investigation.

The potential clinical implications of EV-associated Lp(a) biology are considerable. Lp(a) has recently been reported to promote cardiovascular calcification by inducing the release of calcifying EVs,^59^ and several EV populations decrease following Lp(a) apheresis, although the APO(a) content of these vesicles has not been examined. ^60^ Several studies have reported that statin therapy reduces circulating extracellular vesicle concentrations and alters extracellular vesicle composition, suggesting that lipid-lowering therapy may modulate EV biogenesis and release.^61,62^ Together with our kinetic and imaging findings, these observations support the emerging concept that EVs may represent an important interface between Lp(a) metabolism, vascular inflammation, thrombosis, and calcification.

Taken together, our kinetic, proteomic, imaging, and therapeutic observations support a revised model of human Lp(a) metabolism. Rather than behaving as a metabolically static lipoprotein after secretion, Lp(a) appears to comprise two kinetically distinct components. APOB follows the well-established LDL metabolic pathway governing cholesterol transport and particle clearance, whereas APO(a) resides within a slower-turnover compartment that may undergo delayed assembly, recycling, redistribution, or extracellular vesicle–mediated trafficking before reassociating with APOB-containing particles. This model reconciles the marked discordance between APO(a) and APOB kinetics observed in our study with emerging experimental and therapeutic evidence, while providing a mechanistic framework linking Lp(a) metabolism to extracellular vesicles and bioactive protein networks involved in lipoprotein remodeling, complement activation, and coagulation. Collectively, these findings redefine Lp(a) as a dynamic lipoprotein system rather than a static circulating particle, identify extracellular vesicle–associated pathways as promising therapeutic targets, and establish a foundation for future in vivo kinetic studies designed to directly test this model during Lp(a)-lowering interventions.

## Conclusions

In conclusion, our study provides new insights into the metabolic organization of human Lp(a). By integrating stable isotope kinetics, quantitative proteomics, and high-resolution extracellular vesicle imaging, we demonstrate that APO(a) and APOB exhibit markedly different metabolic behavior despite existing within the same circulating Lp(a) particle. These findings challenge the prevailing assumption that APO(a) and APOB remain metabolically coupled after secretion and instead support a model in which APO(a) resides within a distinct kinetic compartment that may undergo delayed trafficking, recycling, or reassociation with specialized APOB-containing particles. The accompanying proteomic and imaging data further suggest that Lp(a) is a dynamic carrier of bioactive proteins capable of interacting with extracellular vesicles, thereby linking lipoprotein metabolism with inflammatory, thrombotic, and extracellular matrix pathways. Together, these observations provide a revised conceptual framework for Lp(a) biology and establish new mechanistic hypotheses that may inform the development and interpretation of emerging Lp(a)-targeted therapies.

## Study limitations

First, this physiological human kinetic study cannot directly define the intracellular or extracellular mechanisms governing Lp(a) assembly, remodeling, or clearance. Accordingly, the mechanistic models proposed here—including a potential role for extracellular vesicles in APO(a) trafficking or Lp(a) assembly—should be viewed as biologically plausible hypotheses supported by the convergence of our kinetic, proteomic, and imaging data rather than as definitive proof. Second, the proteomic analyses were performed in a relatively small, deeply phenotyped kinetic cohort (n = 16). Although correction for multiple comparisons reduced statistical power and increased the likelihood of Type II error, several nominally associated proteins clustered within well-established pathways of lipoprotein remodeling (CETP, PLTP, APOD) and complement activation (C3) supporting the biological plausibility of these findings while underscoring the need for validation in larger independent cohorts. Third, our kinetic analyses quantified whole-particle Lp(a)-APO(a) and APOB turnover and therefore could not distinguish the metabolic behavior of individual APO(a) isoforms. Finally, although our sample size is comparable to prior stable isotope investigations of Lp(a) metabolism, it limited our ability to detect modest associations and evaluate subgroup differences across diverse populations.

Despite these limitations, the integration of stable isotope tracer kinetics, quantitative proteomics, extracellular vesicle characterization, imaging flow cytometry, and super-resolution microscopy provides, to our knowledge, the most comprehensive multidimensional assessment of human Lp(a) metabolism to date and establishes a framework for future mechanistic studies of Lp(a) assembly, remodeling, and clearance.

## Acknowledgments

We would like to thank our study participants and the CUIMC IICTR CTSA nurses and Bionutrition Unit members. We thank the Columbia Stem Cell Initiative Flow Cytometry Core Facility at CUIMC for assisting with establishing a method to image our EV/apoprotein association and our collaborators at ONI for assisting with optimization of EV imaging study. We thank Drs. Henry Ginsberg for the scientific discussions during protocol development and Dr. Sekhar Ramakrishnan for insights on pool fit and lipoprotein modeling. We thank Dr. Vikash Ali for acquiring the TEM images and staining the EVs.

## Author Contributions

GRS conceived and designed the study. GRS, AM, NM, MK, SS and SK acquired the study data. TT performed modeling of kinetic data. GRS, TT, NM, YL and SS performed the statistical analyses and interpreted the results. HW, MK and SMN provided critical intellectual assistant for EV work. GRS drafted and finalized the manuscript. TT, AM, SM, MI, YL and SS critically revised the manuscript for important intellectual content. All authors reviewed and approved the final manuscript and agreed to be accountable for all aspects of the work.

